# Imaging CRISPR-Edited CAR-T Cell Therapies with Optical and Positron Emission Tomography Reporters

**DOI:** 10.1101/2025.06.06.658308

**Authors:** Rafael Enrique Sanchez-Pupo, John Joseph Kelly, Nourhan Shalaby, Ying Xia, Francisco Manuel Martinez-Santiesteban, Jasmine Lau, Ivy Verriet, Matthew Fox, Justin Hicks, Jonathan Dale Thiessen, John Andrew Ronald

## Abstract

**Rationale:** Chimeric antigen receptor (CAR) T cell therapies have shown remarkable success in treating hematological cancers and are increasingly demonstrating potential for solid tumors. CRISPR-based genome editing offers a promising approach to enhance the potency and safety of CAR-T cells. However, several challenges persist, including inefficient tumor homing and treatment-related toxicities in normal tissues, which continue to hinder widespread adoption. Advanced imaging technologies, including bioluminescence imaging (BLI) and positron emission tomography (PET), provide real-time insights into CAR-T cell distribution and activity in vivo, both in preclinical models and in patients. Here, we developed *Trackable Reporter Adaptable CRISPR-Edited CAR* (tRACE-CAR) T cells, a modular system for site-specific integration of CARs and imaging reporters.

**Methods:** The luciferase reporter AkaLuciferase (AkaLuc) or the human sodium iodide symporter (NIS) were cloned downstream of the CAR in adeno-associated virus (AAV) donors for BLI or PET tracking, respectively. CARs with imaging reporters were knocked into the *TRAC* locus of primary human T cells via CRISPR editing and AAV transduction. Editing efficiency was evaluated by flow cytometry and junction PCR. In vitro cytotoxicity was assessed by BLI using firefly luciferase (Fluc)-expressing cancer cells co-cultured with CAR-T cells at varying effector-to-target ratios. In vivo, BLI and PET imaging assessed CAR-AkaLuc and CAR-NIS T cell expansion and trafficking in Nod-SCID-gamma mice bearing xenograft tumors.

**Results:** T cell receptor (TCR) knockout efficiency exceeded 85%, with CAR expression observed in 70–80% of cells, depending on the reporter used. Reporter-engineered CAR-T cells retained functionality in vitro and exhibited significant cytotoxicity against target cancer cells, outperforming naïve T cells. In vivo, AkaLuc BLI and ^18^F-tetrafluoroborate PET enabled non-invasive tracking of viable CAR-T cells. Notably, the route of administration (intravenous, peritumoral, or intraperitoneal) significantly influenced the distribution of CAR-T cells and their therapeutic effectiveness.

**Conclusion:** tRACE-CAR enabled precise optical and PET tracking of CAR-T cells in models of B cell leukemia and ovarian cancer, allowing dynamic, non-invasive monitoring of cell distribution in both tumors and off-target tissues. This imaging platform could lead to more personalized, effective CRISPR-edited CAR cell therapies.

## Introduction

Chimeric antigen receptors (CAR) are synthetic proteins designed to enhance the antigen-specific recognition and attack of cancer cells by immune cells such as T lymphocytes [1,2]. CAR-T cell therapy is now considered a standard treatment for patients with relapsed or refractory B-cell lymphomas, B-cell acute lymphoblastic leukemia, and multiple myeloma [3,4]. Recent successes in patients with brain cancer (e.g., neuroblastoma) have renewed excitement about the applicability of CAR-T cells for solid tumors [5]. However, significant challenges remain, like patient non-response/relapse and adverse effects such as cytokine release syndrome (CRS), immune effector cell-associated neurotoxicity syndrome (ICANS), and on-target/off-tumor harm of healthy cells [4,6–9]. In solid tumors, poor CAR cell trafficking, antigen heterogeneity, and immunosuppressive microenvironments can also reduce therapy effectiveness [10]. These challenges emphasize the need for new strategies to enhance CAR-T cell potency and mitigate side effects, as well as technologies for objective monitoring of CAR-T cell fate to understand patient outcomes and side effects.

To improve the therapeutic index of CAR-T cells, many groups have used clustered regularly interspaced short palindromic repeats (CRISPR)-based genome editing to precisely knock-out certain endogenous genes, knock-in transgenes of interest at specific loci, or both [11]. In 2017, a landmark study used CRISPR to knock-in an anti-CD19 CAR transgene into the *TRAC* locus of human T cells, disrupting T cell receptor (TCR) assembly and enhancing therapeutic efficacy against B-cell leukemia in mice, compared to lentiviral-engineered cells [12]. Clinical trials using CRISPR-based targeting at the *TRAC* loci and/or other loci aimed at making more efficient, safer, and cost-effective CAR cell products are emerging [13–16].

As novel CAR cells are developed and translated, complementary technologies are essential for improved post-infusion monitoring. Current methods assessing circulating CAR cell levels or their products fail to fully capture their activity within tumors or other tissues [17]. Preclinical and clinical reporter gene imaging technologies can offer non-invasive, spatiotemporal insights into CAR cell migration, proliferation, and persistence after infusion. For instance, orthogonal reporters for bioluminescence imaging (BLI) have allowed for highly sensitive tracking of both CAR cells and cancer cells in individual animals [18,19]. Clinically relevant reporters for positron emission tomography (PET) derived from viruses [20,21], bacteria [22], or humans [17,19,23–28] have demonstrated the ability to visualize lentiviral-engineered CAR-T and CAR-Natural Killer (CAR-NK) cells in various preclinical cancer models, and pioneering clinical studies using a virally-derived PET reporter firmly established the ability to track CAR-T cells in high-grade glioma patients [29,30].

To leverage upon CRISPR-edited CAR cells and BLI/PET reporter-based cell tracking, this research focused on developing a site-directed, modular, and efficient system to produce **T**rackable **R**eporter **A**daptable **C**RISPR-**E**dited **CAR** (tRACE-CAR) T cells. We validated this technology by editing T cells at the *TRAC* locus with different CARs and imaging reporter genes, allowing us to visualize CAR-T cell accumulation at both on-and off-tumor sites in models of B cell leukemia and ovarian cancer. The tRACE-CAR system is an affordable solution that can be adapted to any genomic locus, offering relatively high editing efficiency and the ability to track the edited cells in preclinical models (BLI/PET) and patients (PET). Our system has the potential to improve the safety and effectiveness of CAR-T therapies by providing decisive *in vivo* information on CAR-T fate that can lead to improved understanding of tumor response and off-target effects, leading to next-generation CAR development and ultimately improving patient management and therapeutic outcomes.

## Results

### Efficient TRAC-targeted CRISPR-Cas9 editing of primary human T cells with a CAR and imaging reporter genes

To create tRACE-CAR T cells based on the work of Eyquem et al. [12], we developed adeno-associated virus serotype 6 (AAV6) donor vectors co-expressing a CAR and an imaging reporter gene with homologous arms (HA) targeting the first exon of the *TRAC* locus to disrupt TCRα expression (**Figure 1A**). These AAV6 vectors contained a promoter-less CD19CAR or HER2CAR transgene positioned after a splice-acceptor sequence and a T2A self-cleaving sequence to drive expression from the endogenous *TRAC* promoter (**Figure 1B**). For imaging, the cDNA of eGFP, AkaLuciferase (AkaLuc) for BLI [31] or human sodium iodide symporter (NIS) for PET[32] was cloned after the CAR. Given HA length can influence homology-directed repair efficiency [33], we tested vectors with 200-bp and 600-bp HA lengths and varying AAV6 doses (1x10^5^-1x10^6^ viral genomes (VG) per cell) following nucleofection of activated human T cells with *TRAC*-targeted Cas9-gRNA ribonucleoproteins (**Figure S1A**). High-efficiency editing (>85% GFP+TCR-) was achieved even at the lowest AAV dose, and no notable differences were found between AAVs with different HA lengths (**Figure S1B**). Overall, comparable results were obtained when switching either the CAR or reporter gene, achieving >90% efficiency in knocking out TCR and rendering 70-80% CAR+TCR-cells (**Figure 1C, D and H**). For the CAR-GFP constructs, CAR expression correlated with GFP expression, and CAR-GFP expression did not change the proportions of CD4+ and CD8+ T cell subtypes compared to the parental cells from the same donor (**Figure S1C-E**). AAV6 donor integration was confirmed by junction PCR from genomic DNA (**Figure 1E and I**).

**Figure 1.**
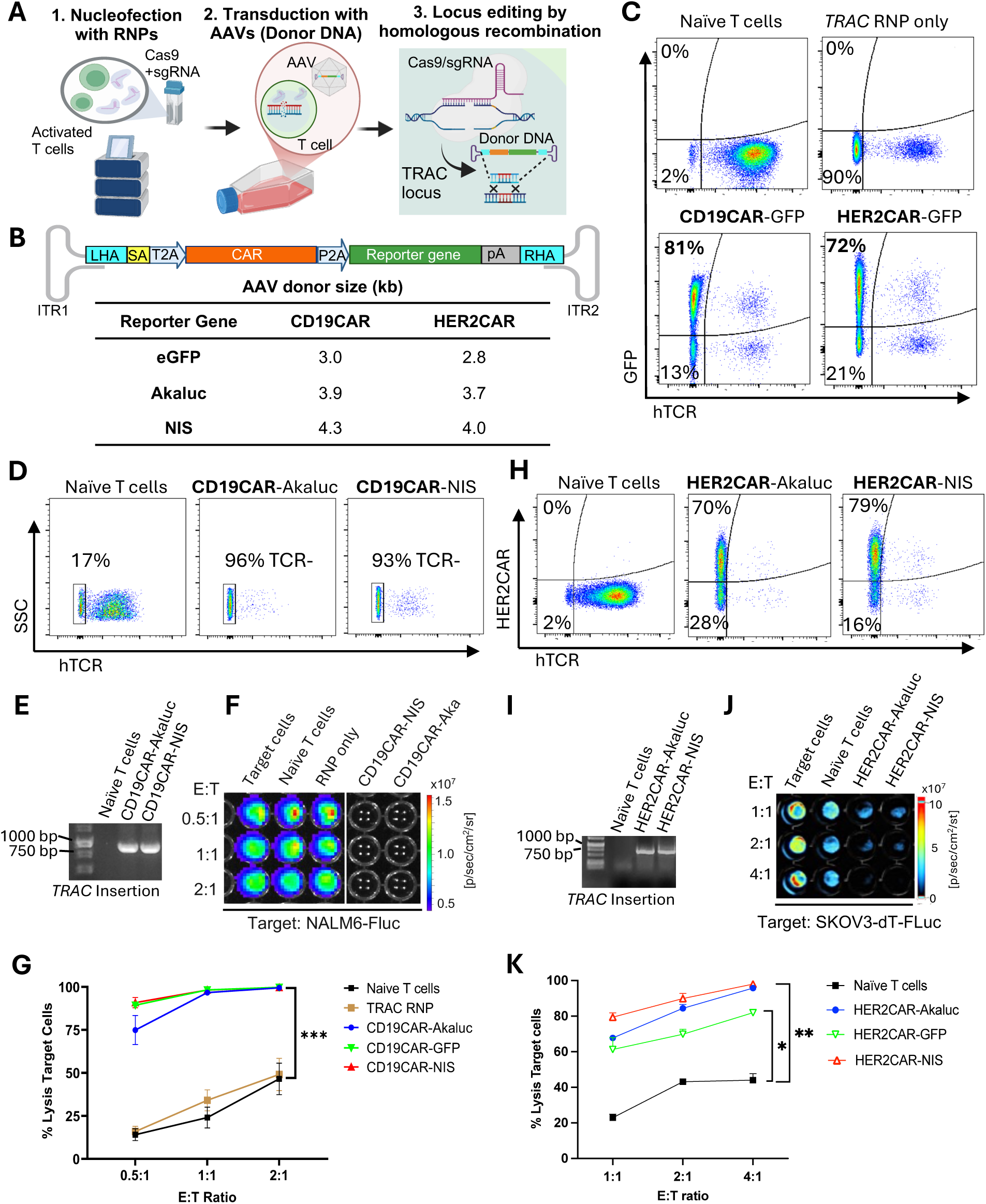
Efficient genome editing of primary human T cells with CAR and imaging reporter genes. **A)** Genome editing workflow using CRISPR-Cas9 technology and adeno-associated viruses (AAV) to engineer human T cells at the *TRAC* locus with CAR and imaging reporter genes for fluorescence (eGFP), bioluminescence (AkaLuc), or PET (NIS). Created in BioRender. Ronald, J. (2022) https://BioRender.com/w24n349. **B)** Schematic of generic donor construct cloned within AAV inverted terminal repeats (ITR). LHA and RHA left and right homologous recombination arms; SA, splice acceptor site; T2A and P2A, self-cleaving peptide sequences; CAR, chimeric antigen receptor; pA, poly A tail. The table shows the full size of each DNA donor. **C, D** and **H)** Flow cytometry of primary T cells labeled with an antibody to the human T cell receptor (hTCR), GFP fluorescence, or a fluorokine against the CAR showing *TRAC* locus editing efficiency. **E)** and **I)** Junction PCR analysis confirming successful CRISPR/Cas9-mediated AAV donor insertion into the *TRAC* locus. BLI-based cytotoxicity assays showing targeted CD19-CAR **(F** and **G**) and HER2-CAR **(J** and **K)** killing at various effector (E; T cell) to target (T; NALM6-Fluc or SKOV3-dT-Fluc cells) ratios 24 hrs after co-culture. Mean comparison using a two-tailed paired T-test. Data is representative of three (N=3) independent experiments. Statistical significance is shown as *, p<0.05; **, p<0.01; ***, p<0.001.

Cytotoxicity assays of CD19-positive leukemia (**Figure 1F and G**) or HER-positive ovarian cancer cells (**Figure 1J and K**) expressing Firefly luciferase showed that T cells engineered with either CAR construct performed well *in vitro*, effectively killing cancer cells regardless of the reporter gene co-expressed. As expected, the number of CAR-AkaLuc T cells correlated with the BLI signal and uptake of the NIS-targeted radiotracer ^18^F-tetrafluoroborate (^18^F-TFB) [34] was 4.89-fold higher in *TRAC*-edited CD19CAR-NIS T cells compared to naïve T cells (**Figure S2**).

### In vivo antitumor efficacy and bioluminescence imaging of tRACE-CAR T cells in leukemia and ovarian models

For *in vivo* evaluation of tRACE-CAR cells, we used a dual BLI system to track both NALM6 cancer cells (Antares) and CAR-T cells (AkaLuc) in individual mice. Mice were injected intravenously (IV) with NALM6 cells followed by an IV dose of 10x10^6^ CD19CAR-T cells or naïve T cells 4 days later (**Figure 2A)**. AkaLuc CAR-T signal, which was mainly localized to the thoracic region and sparsely in the abdomen, decreased 3.8-fold from Day 3 to Day 10, but remained stable thereafter (**Figure 2B and C**). Notably, Antares cancer cell signal was reduced from Days 12 to 25 post-treatment in the CAR-T group compared to naïve T cell and untreated control mice, which resulted in significantly extended survival (**Figure 2D-F**).

**Figure 2.**
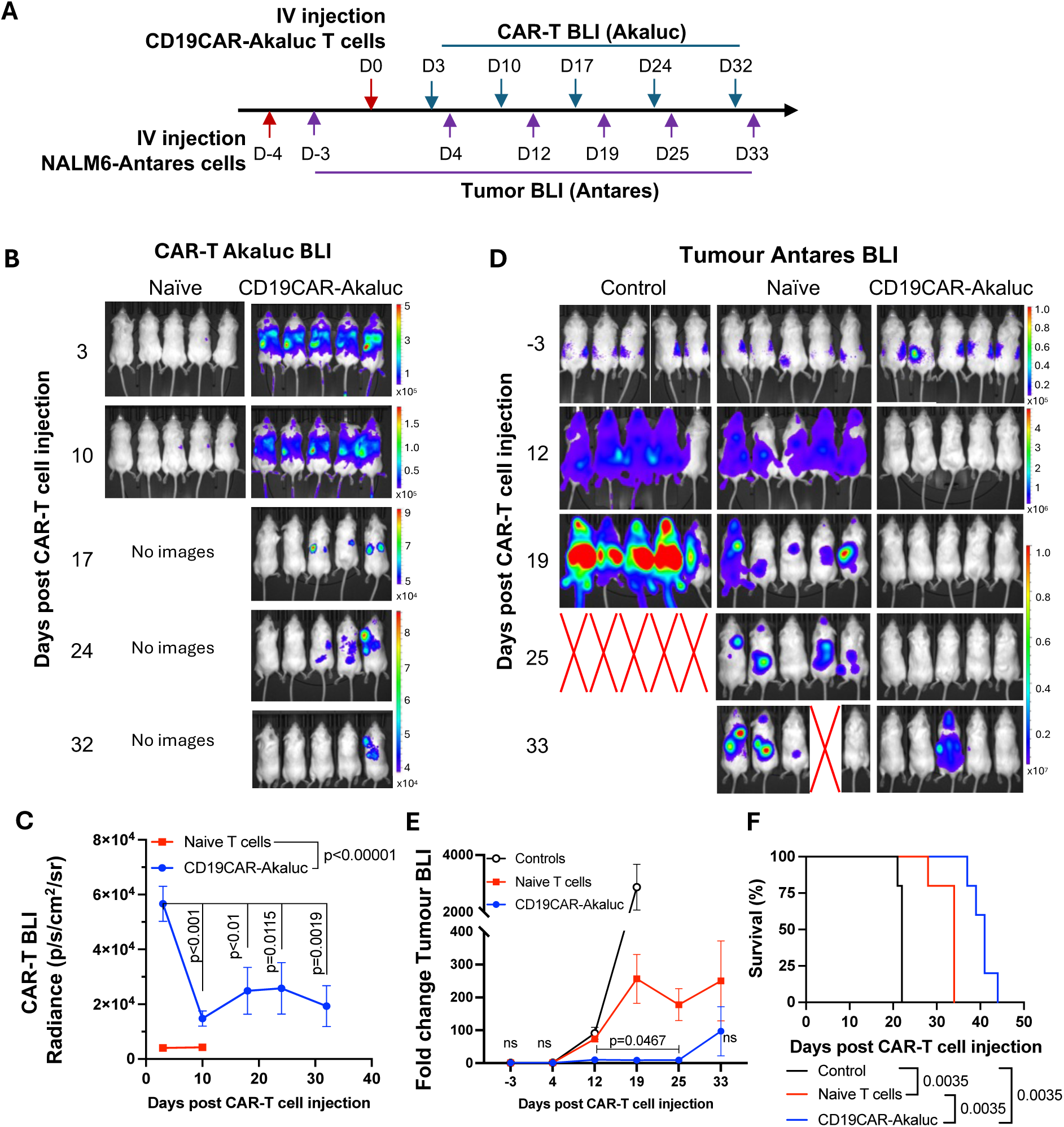
In vivo tracking of human CD19CAR-AkaLuc T cells in a mouse model of NALM6 leukemia. **A)** Experimental timeline of the intravenous (IV) NALM6-Antares leukemia mouse model treated with CD19CAR-AkaLuc T cells (IV administration). **B)** CD19CAR-AkaLuc BLI at various time points, starting 3 days (D3) post CAR-T cell injection. Naïve images were only included up until day 17 for determining the background Akalumine substrate signal. **C)** Quantification of whole-body AkaLuc BLI signal. The difference of CD19CAR-AkaLuc BLI vs background signal in Naïve T cells is shown on the graph for D3 and D10 and was analyzed with an unpaired t-test. Differences in the AkaLuc signal over time starting from D3 were compared with repeated measures ANOVA and Tukey’s multiple comparison test. **D)** Images of NALM6-Antares (tumor) BLI in control (no treatment), naïve T cell, and CD19CAR-AkaLuc T cell treatment. Red X denotes animal death before imaging timepoint. **E)** Quantification of tumor burden based on Antares BLI. Statistical significance is shown from comparisons of Naïve T cell vs CD19CAR-AkaLuc treatments at each time point. Analyzed using a Mann-Whitney test with the Holm-Šídák method for correction for multiple comparisons. **F)** Survival curve (days from post-NALM6-Antares IV infusion). Data was analyzed with a Log-rank test. N = 5 for each group. Data is presented as mean ± SD. Statistical significance is shown in graphs. ns, non-statistically significant.

We then evaluated tracking of AkaLuc-expressing HER2CAR-T cells in mice bearing HER2-positive ovarian SKOV3-ip1 subcutaneous (SC) xenografts (**Figure 3A**). One day following IV infusion of 5x10^6^ HER2CAR-T cells, AkaLuc signal was primarily concentrated in the thoracic region (**Figure 3B and C**). By Day 14, HER2CAR-T cells localized to SC tumors, and expanded over time. A significant increase in AkaLuc CAR-T signal was also detected in the lungs (**Figure 3C**). Near endpoint, mice receiving

**Figure 3.**
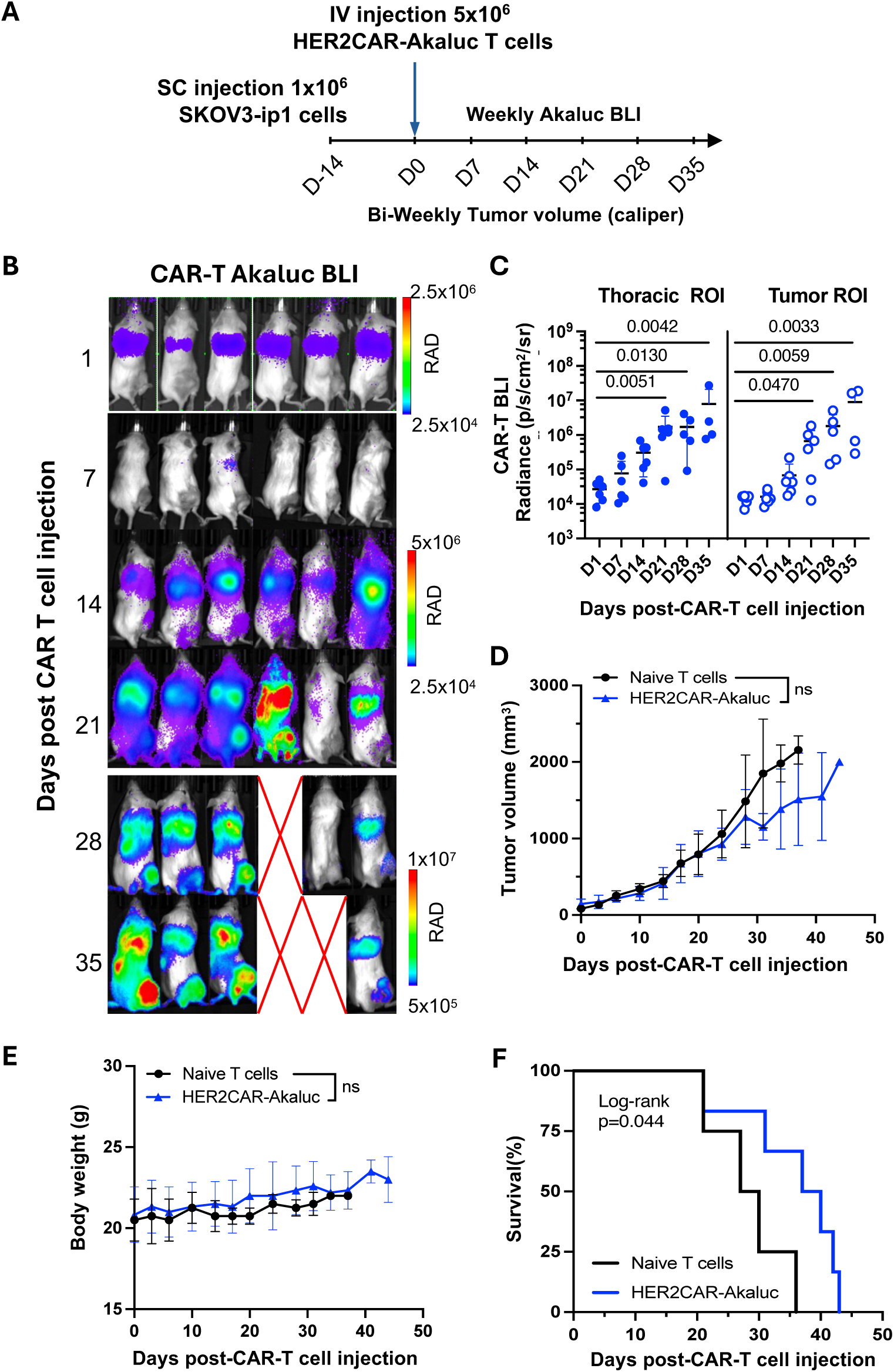
BLI tracking of HER2CAR-AkaLuc T cells in a subcutaneous model of ovarian cancer. **A)** Timeline of the subcutaneous (SC) SKOV3-ip1 ovarian cancer cell model to longitudinally monitor HER2CAR-AkaLuc T cell localization and expansion. **B)** BLI of HER2CAR-AkaLuc treated mice after IV CAR-T cell injection. RAD color scale: average Radiance (p/sec/cm^2^/sr). **C)** Quantification of average radiance from HER2CAR-AkaLuc T cells in thoracic and tumor ROIs. Data was analyzed and compared to the first-day post-CAR-T cell injection using a Kruskal-Wallis test followed by Dunns’s test. **D)** Tumor volume measurements. **E)** Mouse body weight measurements. Data was analyzed using an unpaired t-test with Welch correction. ns, non-statistically significant differences. **F)** Survival curve. Naïve T cell (n=4) vs. HER2CAR-AkaLuc (n=6) mice. Mean ± SD is shown, with symbols indicating individual animal data points.

CAR-T cells showed a trend towards lower tumor volume compared to the naïve T cell group, without changes in body weight between treatment groups (**Figure 3D and E**). Based on tumor volume and other humane endpoints set in our experimental protocol, HER2CAR-T cell treatment significantly improved mouse survival compared to naïve T cells **(Figure 3F**).

Overall, our results show that our tRACE-CAR system allowed for effective BLI of therapeutic cells expressing different CARs in both hematological and solid cancer models.

### [^18^F]-TFB PET tracking of HER2-targeted tRACE-CAR cells in an ovarian cancer subcutaneous xenograft model

We next evaluated the ability to track both on- and off-tumor localization of tRACE-CAR T cells using the sodium iodide symporter (NIS), a clinically translatable, human-derived PET reporter. Mice bearing subcutaneous SKOV3-ip1 tumors expressing Firefly luciferase were randomized to receive IV injections of 5×10⁶ HER2CAR-NIS T cells or saline (**Figure 4**). To potentially enhance therapeutic efficacy, an additional cohort received peritumoral (PERT) injections of CAR T cells, as previously described [35]. [¹⁸F]-tetrafluoroborate ([¹⁸F]-TFB) PET imaging was performed on Days 17 and 28 post-treatment (Day 28 only for saline controls), and tumor BLI was conducted weekly (**Figure 4A**).

**Figure 4.**
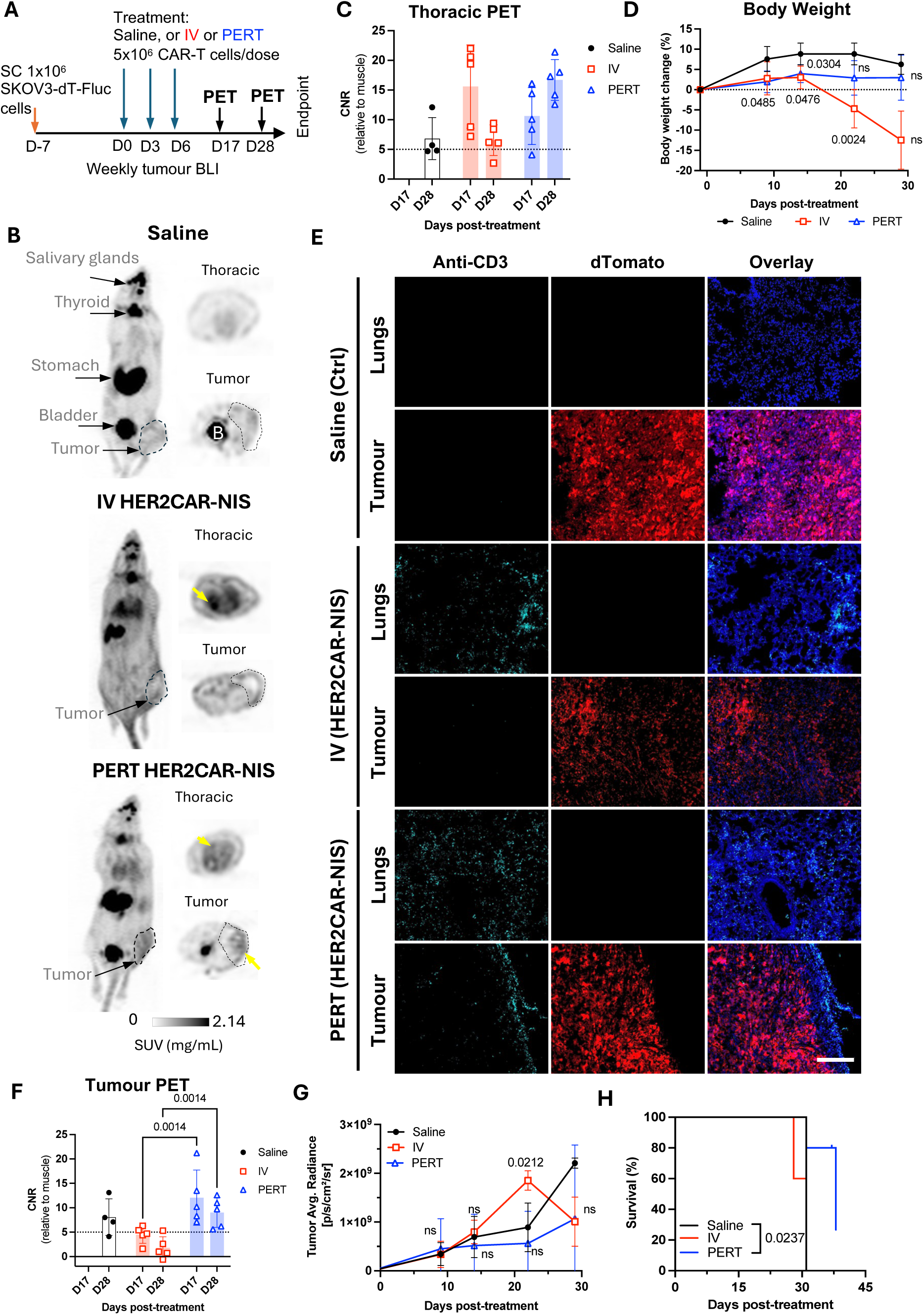
PET imaging and tissue analysis of HER2CAR-NIS T cells in a subcutaneous solid tumor model of ovarian cancer. **A)** Experimental timeline of the SC inoculation of SKOV3-ip1 (dTomato^+^Fluc^+^) tumor cells followed by three doses of IV or peritumoral (PERT) injection of HER2CAR-NIS T cells. **B)** Maximum intensity projections (MIP) showing coronal views and single-slice axial images of representative tumor-bearing mice on Day 28 after treatment. Grayscale bar displays SUV (g/ml) corresponding to the maximum and minimum intensity levels. Yellow arrows point to areas of increased radiotracer uptake. Physiological radiotracer uptake is highlighted in salivary glands, thyroid, stomach, and bladder. Tumor locations are circled with a black dashed line. **C)** and **F)** Comparison of CNR values (relative to muscle) from the volumes of interest (VOIs) of the thoracic and tumor ROI, respectively. Data was analyzed with an ordinary two-way ANOVA followed by a Tukey multiple comparison test. Symbols indicate individual animal data points. **D)** Percent body weight change after CAR-T cell treatment. **E)** Immunofluorescent staining of CD3-positive cells (cyan) in lung and tumor sections derived from the same mice shown in B. Tumor cells expressed dTomato (red) and sections were counterstained with DAPI to label nuclei (blue). Scale bar = 200 µm. **G)** Average radiance of tumors (Fluc BLI) over time. Data was analyzed using a mixed-model ANOVA and Tukey’s multiple comparison test. **H)** Survival curves analyzed with a Log-rank test. p-values indicate statistical significance compared to saline control. At least N=4 mice per group. ns, non-statistically significant.

Consistent with the known biodistribution of [¹⁸F]-TFB [36], saline-treated mice exhibited radiotracer uptake in the thyroid, salivary glands, stomach, and bladder (**Figure 4B**). In contrast, mice treated with HER2CAR-NIS T cells via either IV or PERT injection displayed pronounced radiotracer accumulation in the lungs which was absent in saline-treated controls (**Figure 4B**). Quantitative analysis revealed lung-to-muscle contrast-to-noise ratio (CNR) PET values exceeding 5, the Rose criterion for detectability [37], for most PERT-treated mice on Days 17 and 28, and for IV-treated mice on day 17, but not Day 28 (**Figure 4C**). By day 28, IV-treated mice exhibited signs of morbidity, including significant weight loss (**Figure 4D**), labored breathing, lack of responsiveness, dehydration, and a low body score. Many IV-treated mice also lacked detectable bladder signal, suggesting impaired renal clearance of the radiotracer (**Figure 4B**).

Although PET signal in the lungs remained elevated in these mice (**Figure S3A**), increased signal in muscle (**Figure S3B**) led to reduced lung-to-muscle CNR at Day 28. Notably, PERT-treated mice showed delayed pulmonary accumulation of HER2CAR-NIS T cells, with significantly lower lung signal at day 17 compared to IV-treated animals (**Figure S3A**). Immunofluorescence staining of lung sections from IV- and PERT-treated mice that exhibited high pulmonary PET signal confirmed the presence of human CD3⁺ T cells at endpoint, supporting the imaging findings (**Figure 4E**, lungs panel).

Tumor-to-muscle CNR PET values were comparable between IV-treated and control mice (**Figure 4F**), consistent with the absence of human CD3⁺ T cells in tumor sections from these groups (**Figure 4E**, tumor panel), and a lack of tumor response (**Figure 4G**). In contrast, PERT-treated mice exhibited significantly higher CNR values compared to the IV-treated group at both Days 17 and 28. Correspondingly, CD3⁺ T cells were observed at the tumor periphery in PERT-treated animals. While tumor BLI signal did not differ significantly between groups (**Figure 4G**), 2 out of 5 PERT-treated mice demonstrated a qualitative reduction in tumor burden via BLI (**Figure S3C-D**), which was associated with significantly improved survival compared to saline-treated control mice (**Figure 4H**).

### PET tracking of tRACE-CAR T cells following locoregional administration in an intraperitoneal ovarian cancer model

Although subcutaneous tumor models are informative for early imaging studies, ovarian cancer predominantly disseminates within the intraperitoneal (IP) cavity [38]. To better mimic the clinical context, we next assessed reporter imaging in an experimental IP metastasis model of ovarian cancer. To attempt to minimize CAR T cell entrapment in the lungs, we administered tRACE-CAR T cells via IP injection. Locoregional delivery of CAR T cells has demonstrated therapeutic potential in several tumor types, including ovarian cancer [39–41].

To determine optimal time points for PET imaging in this model, we injected CRISPR-edited HER2CAR T cells expressing AkaLuc into two mice bearing intraperitoneal SKOV3-ip1 tumors engineered to express Antares. We then performed dual BLI over time (**Figure 5A**). Qualitatively, one mouse partially responded to therapy, based on minimal cancer BLI signal increase over time, while the other mouse did not. Both mice consistently showed CAR T cell signals in the abdominal region at all observed time points while one mouse additionally exhibited an increasing signal in the thoracic region from Days 19 to 50. Based on these dynamics, we selected Days 17 (prior to the appearance of thoracic signal) and 28 (when strong abdominal signals were observed in both mice) as the PET imaging time points.

**Figure 5.**
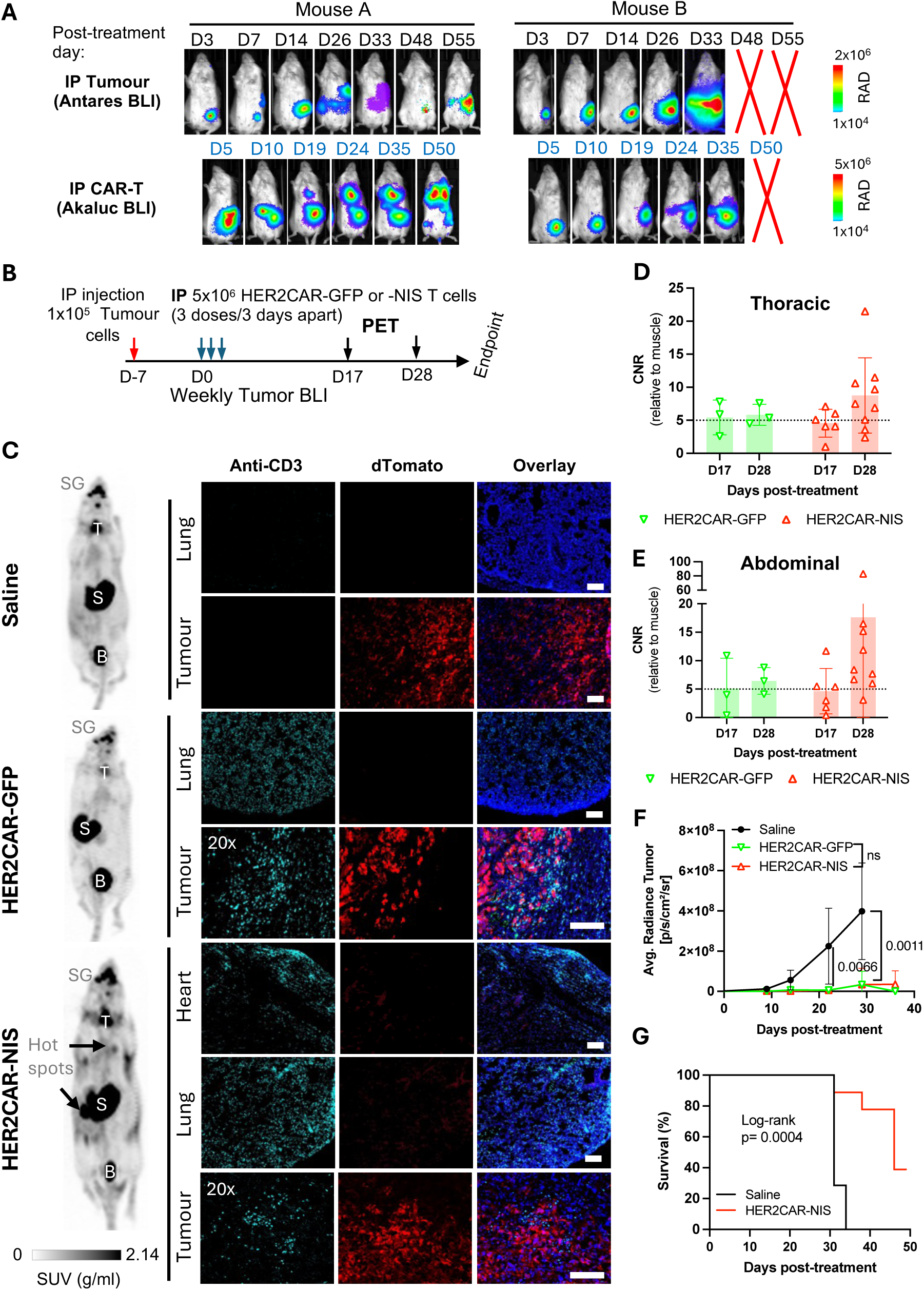
Imaging of HER2CAR T cells after locoregional administration in an orthotopic model of ovarian cancer. **A)** Dual BLI for simultaneous tracking of IP injected HER2CAR-AkaLuc T cells and ovarian SKOV3-ip1-Antares tumor cells within the same mouse. **B)** Timeline of IP SKOV3-ip1(dT^+^Fluc^+^) cancer model with IP-administered HER2CAR-NIS T cells. D17, and D28 refer to PET imaging time points after the first dose of CAR-T cells. **C)** Representative ^18^F-TFB PET maximum intensity projection (MIP) images of tumor-bearing mice treated with saline, HER2CAR-GFP, or HER2CAR-NIS T cells at D28 (left panel). Black arrows highlight hotspot regions of ^18^F-TFB accumulation. Basal tracer uptake in the thyroid (T), salivary glands (SG), stomach (S), and bladder (B) is shown. Anti-CD3 immunofluorescence labeling (cyan) of mouse tissue samples collected from matching treatment groups (right panels). Tumor cells express dTomato (red) and nuclei were counterstained with DAPI (blue). Scale bars = 100 µm. **D)** and **E)** Comparison of the CNR values (relative to muscle) calculated from the entire thoracic and abdominal VOI regions showing an increased CNR (non-statistically significant) in animals treated with HER2CAR-NIS at D28. Data was analyzed with a mix-effect model followed by a Fisher’s LSD test. Symbols indicate individual animal data points. **F)** Average tumor radiance per treatment group showing significant treatment effect in animals treated with HER2CAR-GFP and -NIS T cells compared to saline controls. One-way ANOVA and Tukey’s multiple comparisons tests was done in the log-transformed values at the indicated timepoints. **G)** Survival curve analysis between Saline-and HER2CAR-NIS-treated mice. Analysis performed using a Log-Rank test. Animal number per groups used for PET imaging: Saline (N=4), HER2CAR-GFP (N=3) and HER2CAR-NIS (N=9). Where indicated, statistical significance is shown. ns, non-statistically significant.

For our PET imaging study, mice received three IP injections of either saline (n = 7), HER2CAR-GFP T cells (n = 4), or HER2CAR-NIS T cells (n = 9), with a total dose of 15 × 10⁶ cells per mouse (**Figure 5B**). [¹⁸F]-TFB PET imaging showed similar tracer biodistribution in HER2CAR-GFP and saline-treated mice. In contrast, some HER2CAR-NIS–treated mice exhibited localized tracer accumulation in the lungs (**Figure 5C**).

Quantification of [¹⁸F]-TFB uptake in the thoracic region revealed higher CNR values in several HER2CAR-NIS mice compared to the HER2CAR-GFP group on Day 28 (**Figure 5D**). Importantly, immunostaining identified CD3⁺ T cells in the lungs of both HER2CAR-NIS and HER2CAR-GFP cohorts, indicating that the elevated PET signal observed in the HER2CAR-NIS group is attributable to NIS co-expression.

Some HER2CAR-NIS–treated mice also exhibited focal regions of elevated PET signal within the abdomen (**Figure 5C**), reflected by increased CNR values (**Figure 5E**). CD3⁺ T cells were detected by immunostaining within dTomato⁺ tumors in both HER2CAR-GFP and HER2CAR-NIS cohorts, further supporting the specificity of NIS-mediated PET imaging (**Figure 5C**). BLI revealed tumor signals confined to the abdominal cavity in all mice. Both HER2CAR-GFP and HER2CAR-NIS groups demonstrated significantly reduced tumor burden compared to saline-treated controls, with no significant differences in treatment efficacy between GFP- or NIS-expressing cohorts (**Figure S4** and **Figure 5F**). HER2CAR-GFP mice were euthanized after the final PET timepoint for tissue analysis, whereas HER2CAR-NIS–treated mice were monitored for long-term outcomes and exhibited significantly prolonged survival relative to the saline group (**Figure 5G**).

Together, our results demonstrated that locoregional therapies (PERT or IP) greatly improved tumor response and increased survival rates in mice with ovarian cancer SC and IP xenografts, respectively. More importantly, we demonstrated that tRACE-CAR is an effective method for non-invasive detection of CRISPR-edited CAR-T cells in tumors and non-targeted tissues using a clinically relevant PET reporter.

## Discussion

CAR-T cells have transformed the treatment of hematologic cancers and are now being explored in solid tumors. While CRISPR-mediated editing can enhance CAR-T potency and is under clinical investigation [42], challenges like heterogeneous responses and on-target/off-tumor toxicities persist. These limitations highlight the need for improved, tissue-specific monitoring tools beyond biopsies or peripheral blood, which may not fully reflect *in vivo* dynamics. Here, we developed a CRISPR-Cas9/AAV6-based strategy to site-specifically integrate both CAR and imaging reporter genes into the *TRAC* locus, termed tRACE-CAR. This approach leverages endogenous regulatory control to produce trackable and potentially safer and more potent CAR-T cells compared to conventional viral transduction [12]. Using this platform, we edited T cells with CD19- and HER2-specific CARs together with AkaLuc (BLI) and NIS (PET) reporter genes and evaluated them in leukemia and ovarian cancer models. Whole-body optical and PET imaging revealed inter-animal differences in trafficking, persistence, and expansion across tumors and normal tissues, illustrating how CAR design, tumor type, and delivery route shape therapeutic behavior. These results demonstrate the value of integrated imaging for optimizing next-generation CAR-T cell therapies.

Early reporter gene strategies used zinc-finger nucleases to target the adeno-associated virus integration site 1 (AAVS1) safe-harbor locus, enabling multimodal imaging in stem cell-derived systems [43]. CRISPR-Cas9 has since improved the versatility and efficiency of targeted knock-ins across diverse cell types [44–46]. For example, Lin et al. inserted a constitutive NIS reporter into AAVS1 in iPSC-derived hematopoietic cells to enable PET tracking of implanted cardiomyocytes [46]. We previously applied homology-independent targeted integration (HITI)-based CRISPR editing with minicircles to image small cancer cell populations with BLI and MRI in vivo [47]. However, most approaches relied on single-cell clones, limiting clinical use in primary immune cells. To address this, we implemented a bulk-editing strategy combining Cas9 RNPs with AAV6 donors in human T cells, consistently achieving >85% editing at TRAC and >70% CAR expression. These efficiencies match or exceed recent reports using Cas9 [48], Cas12a [49], or non-viral donors [50], and support scalable, clinically relevant production of imageable CAR-T cells.

BLI is a mainstay for indirect cell tracking in preclinical cancer models due to its ease of use, ability to image more than one animal at a time and being relatively affordable.

Most studies have used BLI to track treatment response, typically tumor clearance, following CAR-T treatment [51–53]. However, these studies can neglect CAR cell trafficking to off-tumor organs, which requires extensive post-sacrifice histological analysis. To address these limitations, recent studies, including our own, have applied dual BLI strategies using orthogonal reporter systems [19,54–56], or spectral unmixing approaches that enable the simultaneous imaging of two distinct luciferases using the same substrate [57]. This provides the opportunity to look beyond just treatment response and offers spatiotemporal whole-body information of CAR cell trafficking. To our knowledge, the current study represents the first use of dual BLI to track CRISPR-edited CAR-T cells in mouse models of blood cancer and solid tumors. Our approach revealed both intratumoral accumulation and off-tumor trafficking, including significant lung localization of HER2CAR-T cells in mice with ovarian tumors, even following intraperitoneal injection.

We showed that targeted integration of CAR and NIS genes at the TRAC locus enables PET imaging of CAR-T cells at low copy numbers (∼1–2 per cell), unlike prior viral approaches that has the potential to randomly integrate multiple copies of NIS [24–26]. While NIS has background uptake in tissues like thyroid and bladder and modest sensitivity (∼10⁴–10⁵ cells), it remains a clinically relevant reporter due to its safety, tracer compatibility, and potential for both imaging and ablation [58]. Future strategies could benefit from PET reporters with alternative tracer biodistributions [59] or antibody-based imaging [60]. Our findings are consistent with previous NIS PET studies tracking CAR-T cells [24–26], including model-dependent persistence in triple negative breast cancer [25] and detection of lung-localized toxicity in a leukemia model [26]. We previously showed that IP-injected HER2CAR-NIS NK-92 cells remained within the peritoneal space in mice with ovarian tumors without toxicity [19], whereas here, HER2CAR-NIS T cells showed lung accumulation, reinforcing that trafficking and toxicity depend on both cell type and delivery route. Our data support IP delivery to enhance efficacy and reduce systemic toxicity, in line with prior ovarian cancer studies [40,41,61,62], and consistent with known HER2CAR-related lung toxicities in mice and a patient [6,63].

Though lentiviral transduction remains the standard [64], recent cases of T-cell malignancies with integrated CAR transgenes have raised concerns [65,66], prompting an FDA boxed warning in 2023 [67]. While causality is unclear [68–70], the risk of insertional mutagenesis with viral vectors [71,72] highlights the appeal of targeted CRISPR-based approaches. Editing at the *TRAC* locus can reduce tonic signaling and exhaustion [12,73,74], and supports universal CAR-T strategies via multiplexed editing of *TRAC*, *PDCD1* (PD-1), *B2M*, and HLA [75–79].

We also observed variability in CAR-T cell expansion and trafficking, as reported in prior studies [17]. Future work could combine imaging with CAR-T phenotyping and tumor profiling (e.g., stemness [80], antigen density [81]) to better understand these dynamics. While we did not perform peripheral blood monitoring, integrating blood data, flow cytometry, and imaging may enhance therapeutic assessment.

In summary, our approach enables precise integration of CAR and imaging reporters into the *TRAC* locus, allowing non-invasive tracking of CAR-T cells in vivo. The tRACE-CAR platform supports broader applications, including multiplex editing or universal CAR-T development [79]. These findings build on earlier PET imaging work in clinical T cell therapy [29] and open new paths for monitoring CRISPR-edited cell therapies in cancer, infection [82], and autoimmunity [83].

## Material and Methods

### Cell lines and culture

NALM6 cells (ATCC: clone G5, #CRL-3273) were grown in RPMI-1640 Medium supplemented with 10% fetal bovine serum (FBS) and 1% antibiotic-antimycotic (A/A) (Thermo Fisher Scientific). HER2-expressing SKOV3.ip1 (RRID:CVCL_0C84) ovarian cancer cells (kindly provided by Dr. Trevor Shepherd, London Regional Cancer Program, University of Western Ontario, ON) were cultured in McCoy’s 5A medium (Thermo Fisher Scientific) containing 10% FBS and 1% A/A. Cells were transduced as previously reported [84] with home-made lentiviral vectors to express dTomato (dT) and Firefly luciferase (NALM6-dt-Fluc and SKOV3-dT-FLuc) or zsGreen and Antares (SKOV3-zsG-Antares). Upon transduction, positive cell populations were sorted based on the fluorescence of dT, zsG, or Antares with a BD FACSAria III cell sorter (BD Biosciences) in the London Regional Flow Cytometry Facility at Robarts Research Institute, University of Western Ontario. All cells were grown in a humidified incubator at 37°C and 5% CO_2_ and were routinely tested for mycoplasma contamination with the MycoAlert PLUS Mycoplasma Detection Kit (Lonza).

### T cell activation and culture

Human peripheral blood mononuclear cells (PBMCs) (STEMCELL Technologies, Cat# 70025.1) were stimulated with Dynabeads Human T-Activator CD3/CD28 (Thermo Fisher Scientific) at a bead-to-cell ratio of 1:1. After 3 days, Dynabeads were removed using an EasySep magnet (STEMCELL Technologies). Activated T cells were cultured in ImmunoCult-XF T cell expansion medium (STEMCELL Technologies) containing 1% A/A and 200 U/mL rhIL-2 (kindly provided by the BRB Preclinical Biologics Repository, Frederick National Laboratory for Cancer Research, USA). The cell culture medium was changed every 2-3 days and cells were maintained at a density of 1 x 10^6^ cells/mL in a humidified incubator at 37°C, 5% CO_2_.

### Adeno-associated virus vector design and production

Adeno-associated virus donor vectors (pAAV) containing AAV serotype 2 inverted terminal repeats were designed so that transgene expression was driven off the endogenous *TRAC* promoter by adding a splice acceptor site (SA) and a self-cleaving T2A peptide in frame with the *TRAC* locus, as described by [12]. A CAR-Imaging reporter gene cassette was cloned immediately after the T2A cleavage sequence, flanked by 200 bp or 600 bp homologous arms (HA sequence) specific to the first exon of the *TRAC* gene. The donor cassette comprised a cDNA encoding a CD19-specific CAR (herein referred to as CD19CAR, kindly provided by Dr’s Holt and Nelson, University of Victoria, Victoria, BC, Canada [85]) or a HER2-targeting designed ankyrin repeat protein (DARPin)-based CAR (referred to as HER2CAR, kindly provided by Dr. Bramson, McMaster University, Hamilton, ON, Canada[63]) followed by a self-cleaving P2A peptide in frame with an imaging reporter gene and the bovine growth hormone (bGH) polyadenylation signal. Specifically, the imaging reporter genes cloned herein were: eGFP, for fluorescence, AkaLuc [86] for bioluminescence imaging (BLI), or the human sodium iodide symporter (NIS) [26] for PET. AAV serotype 6 (AAV6) particles were produced and purified by SignaGen Laboratories (Frederick, MD, USA) and VectorBuilder (Chicago, IL, USA).

### CRISPR/Cas9 and AAV transduction

Activated primary human T cells were nucleofected with ribonucleoprotein (RNP) complexes consisting of Alt-R® S.p. HiFi Cas9 Nuclease V3 (IDT, Cat#1081061) and synthetic *TRAC* sgRNA (5′-CAGGGUUCUGGAUAUCUGU-3′) (Synthego, USA) as per [12] using the P3 Primary Cell 4D-Nucleofector X Kit S (Lonza). To make the RNP complexes, 1 μL of sgRNA (60 μM) and 1 μL of HiFi Cas9 Nuclease (20 μM) were mixed in a 0.2 mL PCR tube (3:1 gRNA:Cas9 molar ratio) and incubated for 10 minutes at room temperature. 1 x 10^6^ activated T cells were pelleted and resuspended in 20 μL of P3 Primary Cell solution + Supplement 1 (Lonza) and mixed with the RNP complexes. Cells and RNPs were transferred to a 16-well Nucleocuvette Strip and nucleofected in a 4D-Nucleofector™ X Unit (Lonza) using the pre-set program E0-115 for stimulated human T cells. Following nucleofection, the cells were left to incubate at room temperature for 5 minutes before adding 80 μL of warm T cell culture medium and incubating for 10 minutes at 37°C, 5% CO_2_. Finally, the cells were gently resuspended and transferred to a 48-well culture plate with 400 μL of the same culture media. Thirty minutes after nucleofection, rAAV6s were added to the cell culture at a multiplicity of infection of 1 × 10^5^ virions per cell (unless described otherwise).

### PCR integration assays

DNeasy Blood and Tissue Kit (QIAGEN) was used to extract genomic DNA from edited T cells following the manufacturer’s instructions. 150 ng of genomic DNA isolated from engineered T cells was used as the template for PCR. A reverse primer was designed to target intron 1 in the *TRAC* locus (5’-TGACTGCGTGAGACTGACTT-3’), and a forward primer in the bovine growth hormone (bGH) polyadenylation signal of the AAV donor cassette (5’-TGGGAAGACAATAGCAGGCA-3’).

### Flow cytometry

Genome-editing efficiency was assessed by flow cytometry using a FACSCanto Cytometer (BD Biosciences) 7-9 days after nucleofection and rAAV6 transduction. Where applicable, rhCD19-Fc-Chimera-Alexa-Fluor® 647 (Bio-Techne, Cat# AFR9269) or rhErbB2/Her2-Fc-His-Alexa Fluor® 647 (Bio-Techne, Cat#: AFR1129) fluorokines were used to measure cell surface expression of CD19CAR or HER2CAR, respectively. Anti-human TCRα/β Phycoerythrin (PE)-Cyanine 7 -conjugated antibody (BioLegend, Cat# 306719) was used to assess TCR expression. Anti-human CD4 PE/Cyanine7 (BioLegend, Cat# 357409) and anti-human CD8a Allophycocyanin (BioLegend, Cat# 300911) were used to assess CD4 and CD8 expression. SYTOX blue^TM^ (Thermo Fisher Scientific) was used as a viability dye to exclude dead cells.

### In vitro BLI and cytotoxicity assays

CAR-T cell cytotoxicity was assessed by co-incubating CAR-T cells with firefly luciferase (Fluc)-expressing target cancer cells and measuring BLI over time. Briefly, 1.25 x 10^4^ target cells/well were plated in 96-well plates in 100 μL of serum-supplemented culture media. T cells were then added at various effector-to-target ratios (0.5:1, 1:1, 2:1, or 4:1). The plate was incubated for 18-24 hours at 37°C, 5% CO_2_. BLI was acquired with an IVIS Lumina XRMS In Vivo Imaging System (PerkinElmer) after adding D-luciferin (150 μg/mL, Syd Labs) to each well. Regions of interest (ROI) were drawn over each well to quantify the peak average radiance per well (photons(p)/second(s)/cm^2^/steradian(sr)). BLI data analysis was conducted using the Living Image Software (PerkinElmer). Percent target cell lysis was calculated using Equation 1:

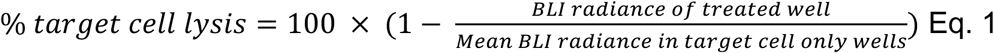

To assess for AkaLuc reporter gene functionality, naïve or AAV-only control (AAV6-CAR-AkaLuc transduction but no CRISPR/Cas9-editing) and CAR-AkaLuc (TRAC-edited + AAV6-CAR-AkaLuc) T cells were seeded in triplicate in 96-well culture black plates and serially diluted from 2, 1, 0.5, 0.25, 0.1, 0.05 (× 10^6) cells/well. Fifteen minutes of BLI acquisition at 37°C was conducted after adding AkaLumine-HCl (TokeOni; Sigma-Aldrich) substrate (0.125 mmol/L, final concentration) to the T cell culture. Peak average radiance was determined from ROIs of each well and quantified as above.

### Animals

All animal experiments received approval from the Animal Care Committee at the University of Western Ontario. 8–13-week-old male and female NOD-scid IL2Rgamma-null (NSG) mice from an in-house colony (Dr. David Hess; Western University) were utilized, except in cases involving SKOV3-derived tumors, where only female mice were employed.

### In vivo Bioluminescence Imaging and Analysis

Mice were anesthetized with 1–2% isoflurane before imaging, and image acquisition was done immediately after intraperitoneal injection of BLI substrates in an IVIS Lumina XRMS In Vivo Imaging System (PerkinElmer). Images were acquired every 2 minutes after substrate injection for up to 30 minutes or until peak radiance (p/s/cm^2^/sr) was reached. The volumes of BLI substrate injected per ∼20 g mouse were 100 μL of Akalumine-HCl (5 mM) (Sigma-Aldrich, Cat#808350-5MG), 100 μL D-luciferin (150 µg/mL) (Syd Labs, Cat#MB000102-R70170), or 50 μL of 2.3 mM Nano-Glo (fluorofurimazine (FFZ), Promega, Cat#N4100) to track AkaLuc-, FLuc- or Antares- expressing cells, respectively. BLI data analysis and visualization were conducted using the Living Image Software (PerkinElmer) and Aura Imaging Software (ver. 4.0.7, Spectral Instruments Imaging).

### Tumor implantation models

For NALM6 leukemia models, 0.5 x 10^6^ NALM6-Antares cells were injected into 6–8-week-old NSG mice via the tail vein and BLI performed the next day to confirm successful injection. Three or four days later, 10^7^ naïve or CD19CAR-AkaLuc cells were injected via the tail vein. BLI of tumors and/or CAR-T cells was then performed at several time points, as indicated in figure timelines.

For subcutaneous models of ovarian cancer, 1 x 10^6^ SKOV3-ip1 (naïve) or SKOV3-ip1-zsG-Antares cells were mixed with Matrigel (50:50 v/v) and injected in the right hind flank. Where applicable, 7-14 days post-tumor implantation, when the tumors reached approximately 100 mm^3^, mice were randomized and received a one-time intravenous injection of 5 x 10^6^ Naïve or HER2CAR-AkaLuc T cells. Caliper measurements were taken, and tumor volume was calculated using the modified ellipsoid formula V = (W(×2) × L)/2, where V is tumor volume, L and W are the longest (length) and shortest (width) tumor diameters, respectively. BLI signal was monitored weekly. When the dual AkaLuc (CAR-T) and Antares (cancer cells) cell-tracking system was evaluated, image acquisition was done 1-2 days apart to avoid background from any residual BLI signal from either BLI reporter gene. Endpoint was considered when the tumor volume reached 2.0 cm^3^, tumors showed signs of ulceration, or the animals exhibited poor health conditions or 20% weight loss. Euthanasia was carried out with isoflurane overdose.

In the intraperitoneal ovarian cancer model, on Day 0, 1 x 10^5^ SKOV3-dT-Fluc cells were suspended in sterile PBS and injected intraperitoneally. Following tumor implantation, three consecutive intraperitoneal doses of saline, 5 x 10^6^ Naïve, or HER2CAR-GFP or -NIS T cells were administered on Days 7, 10, and 13. Tumor burden was evaluated biweekly using BLI. PET imaging was conducted 17 and 28 days after CAR-T cell injection. HER2CAR-GFP-treated mice were used as controls for imaging and were sacrificed for tissue collection before reaching the endpoint; therefore, they were not included in the survival analysis. The endpoint for this tumor model was reached when the animals showed signs of significant tumor growth (tumor radiance ≥8 x 10^8^ p/s/cm^2^/sr), severe abdominal distention due to ascites formation and internal hemorrhage, labored breathing or a 20% weight loss. In such cases, euthanasia was performed using an overdose of isoflurane.

### In vitro [^18^F]-TFB uptake

[^18^F]TFB was synthesized as described previously [87] . Naïve or CAR-NIS-expressing T cells were seeded in a 24-well dish (1 × 10^6^ cells/well), and 0.1 MBq of [^18^F]-TFB was added to each well. Cells were incubated for 45 minutes before centrifuging and washing the cell pellets with ice-cold PBS twice. The activity of harvested cells was measured using a gamma counter, and data was presented as net counts.

### In Vivo [^18^F]-TFB PET Imaging and Analysis

Mice were anesthetized with 2% isoflurane, injected with 10.71 ± 0.68 MBq of [^18^F]-TFB in 50–150 𝜇L sterile saline solution via the tail vein, and imaged with a Siemens Inveon® small animal PET system (Siemens Medical Solutions Inc., USA). Static PET scans were acquired 30 minutes after injection of the radiotracer. Animal breathing rate and body temperature were monitored and maintained between 40-70 bpm at 37 ℃.

Static PET data were acquired for 15 min, and whole-body images were reconstructed using a three-dimensional ordered subsets expectation-maximization (3D-OSEM) algorithm. The radiotracer dose was corrected in relation to the acquisition time and injection time. Quantification of PET signal was performed by manual segmentation of ROIs across coronal slices in each animal to generate volumes of interest (VOIs) using Horos Project software (v4.0.0.RC5). A blinded investigator measured SUV values in muscle (left hind limb), subcutaneous tumors, and thoracic and abdominal cavities (to consider intraperitoneal tumors and CAR-T cell accumulation). Tissues/organs with known [^18^F]-TFB uptake or linked to its renal excretion (e.g., thyroid, stomach, kidneys, bladder) were excluded during ROI segmentation, using PET images from saline control animals as reference.

SUV was calculated with the equation 2:

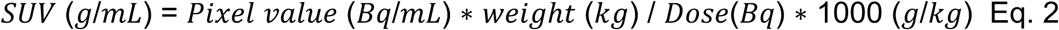

Contrast-to-noise ratio (CNR) was calculated with the equation 3:

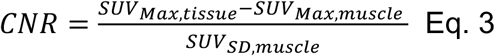

### Histology and Immunofluorescence Staining

Organ tissues and tumors were collected after sacrifice and were fixed overnight in 4% paraformaldehyde (PFA) in PBS at 4°C. Following tissue processing, paraffine-embedding and H&E staining were performed at the Molecular Pathology Core Facility at Robarts Research Institute, Western University. For immunofluorescence staining, 4% PFA-fixed specimens were sequentially incubated overnight in 10%, 20%, and 30% sucrose solutions (in 1X PBS) before OCT embedding and cryo-sectioning. 10 mm-thick frozen sections were thawed and permeabilized with 0.25% (v/v) Triton X-100 (Sigma) in PBS for 10 min and blocked for 1 h with 5 % normal goat serum, and 1 % BSA in PBS at room temperature. Next, tissue sections were incubated overnight at 4°C with rabbit recombinant Anti-CD3 epsilon antibody [EP449E] (Abcam, Cat# ab52959, Cambridge, MA, USA) primary antibody (1:150 dilution; 1.5 μg/mL working concentration). After three consecutive washes with PBS, 5 min each, a secondary Goat anti-Rabbit IgG (H+L) Cross-Adsorbed Alexa Fluor™ 647 secondary antibody (Invitrogen, Cat# A-21244, Eugene, OR, USA) (1:400 dilution; 3.2-μg/ml working concentration) was applied for 1h at RT. Sections were counterstained with Mounting Medium containing DAPI (Abcam, Cat# ab104139, Cambridge, MA, USA). Slides were imaged with 4x/NA 0.13 PhL or 10x/NA 0.30 Ph1 Olympus UPlanFL N objectives in a Revolve-2 Microscope (ECHO, San Diego, CA).

### Statistics

All data are expressed as mean ± standard deviation (SD) of at least three independent experiments or animal subjects. A p-value<0.05 was considered statistically significant. Sample size and the statistical tests are described in the figure legends. Statistical analysis was performed with GraphPad Prism Software (Version 10.0.0 (131), for Mac OS X, GraphPad Software Inc., La Jolla California USA, www.graphpad.com).

## Data availability

The data that support the findings of this study are available from the corresponding author upon reasonable request.

## Abbreviations

AAV: adeno-associated virus; AAV6: adeno-associated virus serotype 6; AAVS1: adeno-associated virus integration site 1; B2M: beta-2 microglobulin; BLI: bioluminescence imaging; CAR: chimeric antigen receptor; CD: cluster of differentiation; CNR: contrast-to-noise ratio; CRISPR: clustered regularly interspaced short palindromic repeats; CRS: cytokine release syndrome; HA: homologous arm; HER2: human epidermal growth factor receptor 2; HITI: homology-independent targeted integration; HLA: human leukocyte antigen; ICANS: immune effector cell-associated neurotoxicity syndrome; IP: intraperitoneal; IV: intravenous; MRI: magnetic resonance imaging; NALM6: human pre-B acute lymphoblastic leukemia cell line; NIS: sodium iodide symporter; NK: natural killer; NSG: NOD scid gamma; PBMC: peripheral blood mononuclear cell; PCR: polymerase chain reaction; PD-1: Programmed cell death protein 1; PERT: peritumoral; PET: positron emission tomography; ROI: Region of interest; RNP: ribonucleoprotein; SC: subcutaneous; SKOV3: human ovarian carcinoma cell line; SUV: standardized uptake value; T2A: Thosea asigna virus 2A peptide; TCR: T cell receptor; [^18^F]-TFB: ^18^F-tetrafluoroborate; TRAC: T cell receptor alpha constant; VOI: volume of interest.

## Authors Contributions

R.E.S-P. and J.J.K. conducted most of the in vitro and in vivo experiments, performed the imaging experiments, analyzed the data, and drafted the manuscript. N.S. assisted with the initial in vitro [^18^F]-TFB uptake assays and PET experiments. Y.X. performed lentiviral transduction and the creation of cancer cell lines. I.V. assisted with CAR-T cell engineering and in vitro killing assays. J.L. assisted with PET image quantification and analysis. F.M.M-S. assisted with the technical setup for mouse monitoring and PET imaging, as well as image analysis. J.H. contributed to the radiotracer synthesis. M.F. and J.D.T. assisted with PET instrumentation, whole-body image acquisition, and reconstruction. J.J.K. and J.A.R. managed the project. J.A.R. secured funding, designed the research, offered analytical guidance, and revised the manuscript incorporating feedback from all authors.

