## Supplemental Figures for "Imaging CRISPR-Edited CAR-T Cell Therapies with Optical and Positron Emission Tomography Reporters"

**Figure S1**

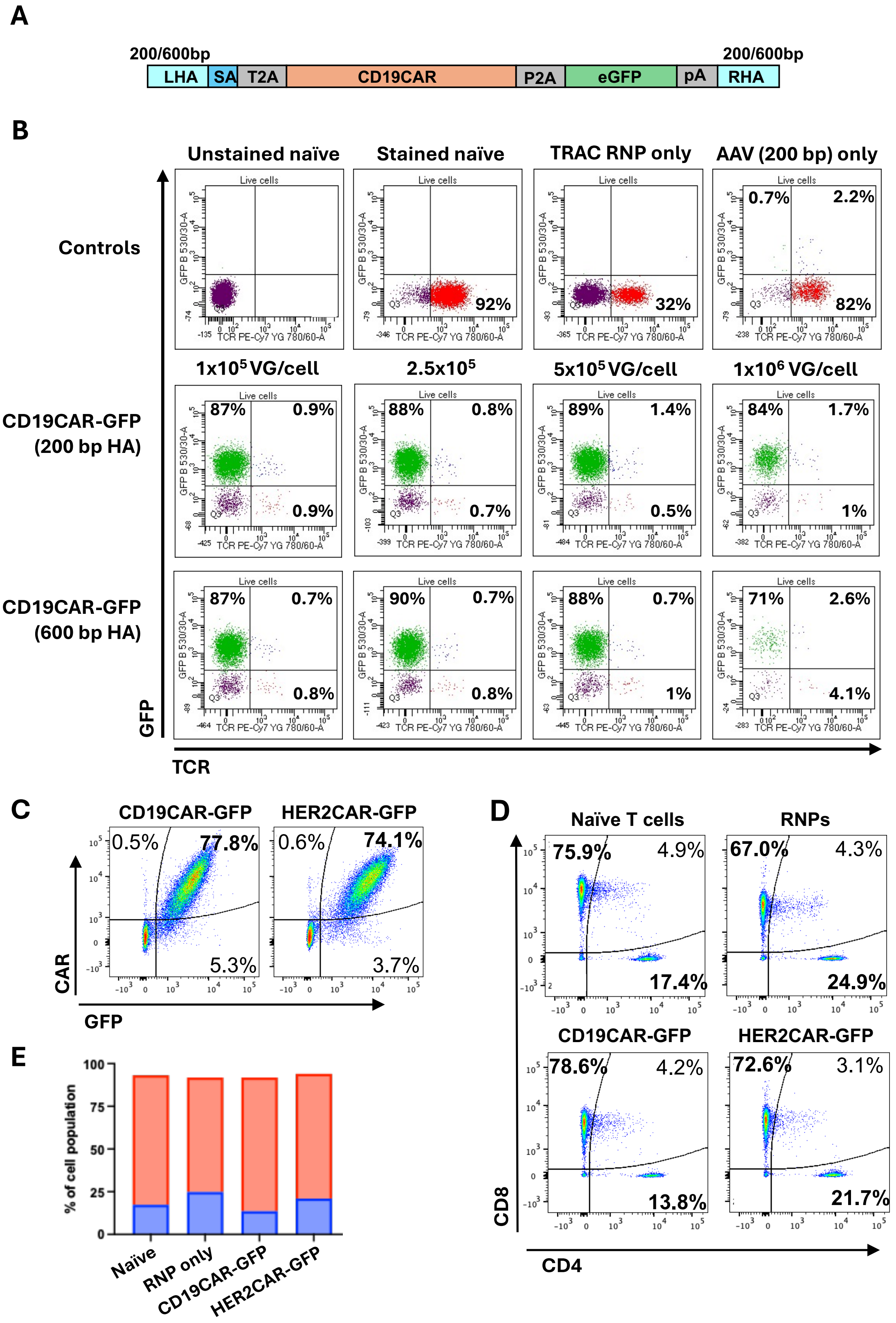
A
200/600bp
LHA
SA
T2A
CD19CAR
P2A
eGFP
pA
RHA
200/600bp
B
Unstained naïve
Stained naïve
TRAC RNP only
AAV (200 bp) only
Controls
92%
32%
0.7%
2.2%
82%
1x10^5 VG/cell
2.5x10^5
5x10^5 VG/cell
1x10^6 VG/cell
CD19CAR-GFP
(200 bp HA)
87%
0.9%
0.9%
88%
0.8%
0.7%
89%
1.4%
0.5%
84%
1.7%
1%
CD19CAR-GFP
(600 bp HA)
87%
0.7%
0.8%
90%
0.7%
0.8%
88%
0.7%
1%
71%
2.6%
4.1%
Live cells
GFP B 530/30-A
TCR PE-Cy7 YG 780/60-A
GFP
TCR
C
CD19CAR-GFP
HER2CAR-GFP
0.5%
77.8%
5.3%
0.6%
74.1%
3.7%
CAR
GFP
D
Naïve T cells
RNPs
75.9%
4.9%
17.4%
67.0%
4.3%
24.9%
CD19CAR-GFP
HER2CAR-GFP
78.6%
4.2%
13.8%
72.6%
3.1%
21.7%
CD8
CD4
E
% of cell population
100
75
50
25
0
Naïve
RNP only
CD19CAR-GFP
HER2CAR-GFP

**Figure S1. *TRAC*-editing efficiency and confirmation of CAR expression in human T cells.**

**A)** Schematic of CD19CAR-GFP donor construct with either 200 bp or 600 bp homologous arms. **B)** Flow cytometry analysis of *TRAC* locus editing efficiency with CRISPR-Cas9 and AAV6 carrying CD19CAR-GFP donors. Activated human T cells were exposed to various amounts of AAV. 1 x $10^6$ AAV genomes/cell caused significant cell death ($4^{th}$ column), hence the lower data outputs. **C)** Flow cytometry analysis of editing efficiency and validation of CAR (labelled with a fluorokine) and GFP co-expression in primary human T cells. **D)** and **E)** Flow cytometry analysis and quantification confirming that CD19CAR- or HER2CAR-GFP knock-in at the *TRAC* locus does not affect the ratio of CD4 to CD8 T cells from the original PBMC donor.

**Figure S2.**

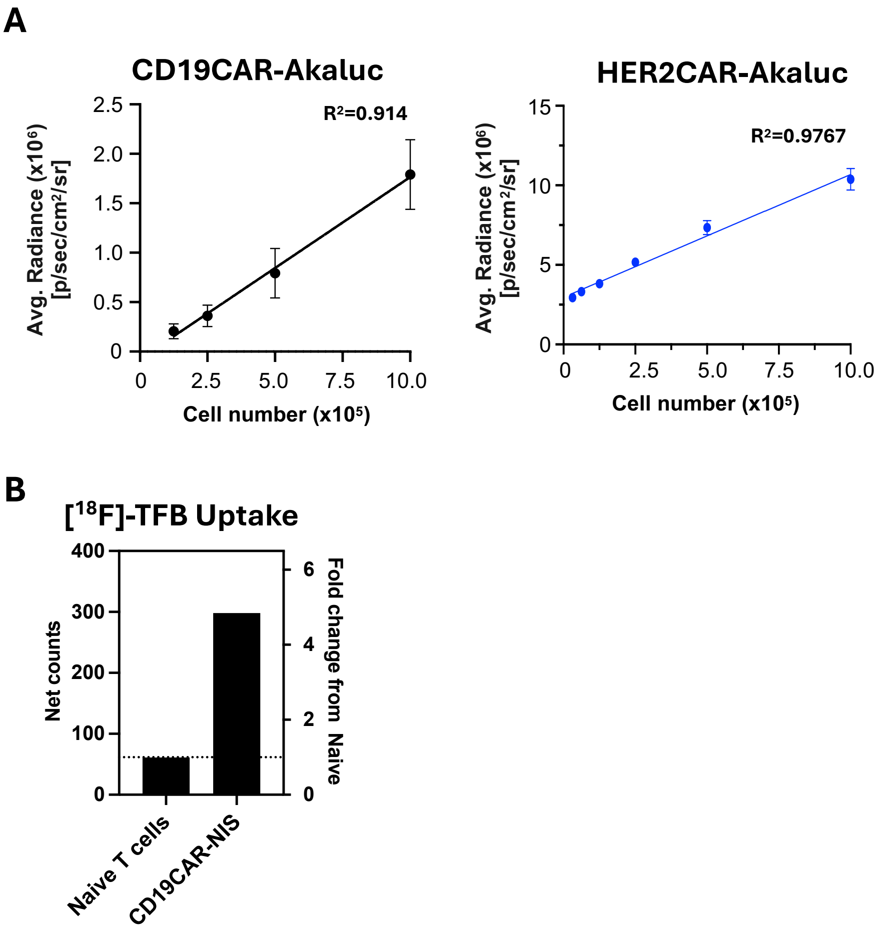


**Figure S2. Functional characterization of imaging reporter genes knocked-in at the *TRAC*-locus.**

**A)** CAR-Akaluc BLI signal correlates with cell number (n = 3). **B)** Gamma counter net counts and fold change from naïve cells of [18]F-TFB uptake into CAR-NIS-expressing T cells.

**Figure S3.**

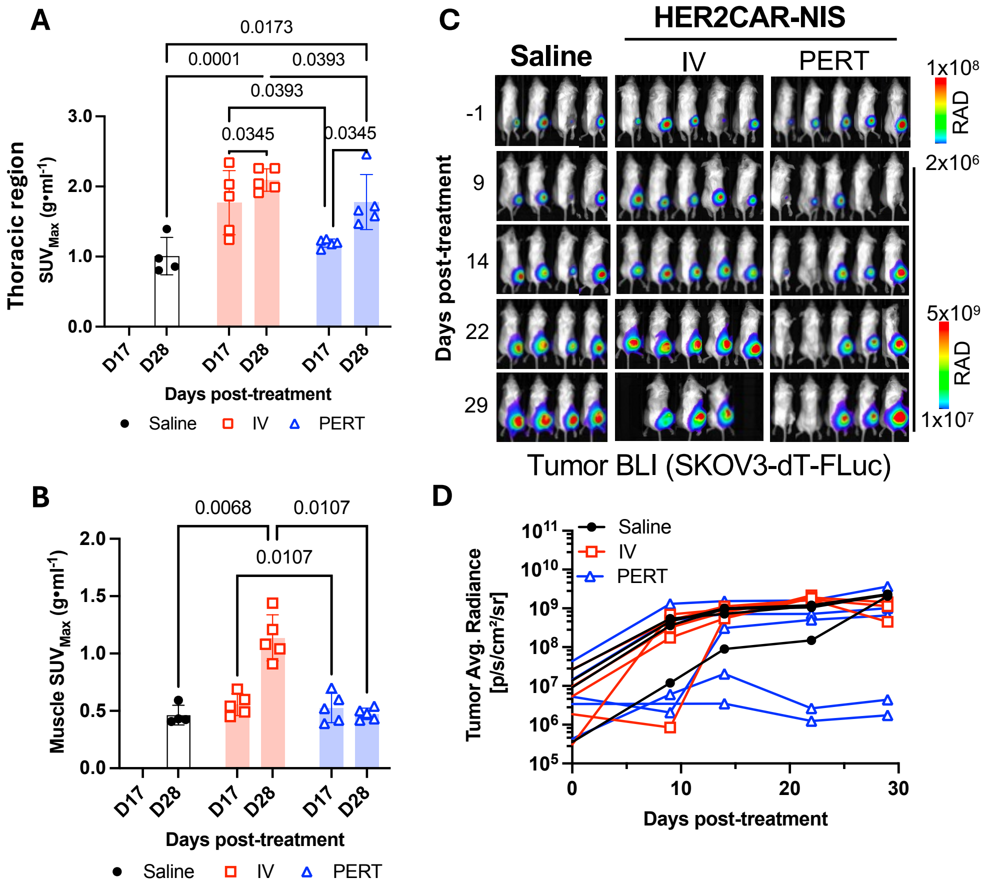


**Figure S3.** ***In vivo*** **[18]F-TFB PET and monitoring of subcutaneous ovarian cancer tumours after HER2CAR-NIS T cell therapy.**

**A-B)** Comparison of Maximum Standardized Uptake Value (SUVMax) from volumes of interest (VOIs) in thoracic and muscle (right thigh) regions of the same animals as in Figure 4. Data was analyzed with an ordinary two-way ANOVA followed by a Tukey multiple comparison test. Symbols indicate individual animal data points. **C-D)** BLI and spaghetti plots of subcutaneous SKOV3-ip1(td+Fluc+) tumour burden after IV or PERT HER2CAR-NIS injections.

**Figure S4.**

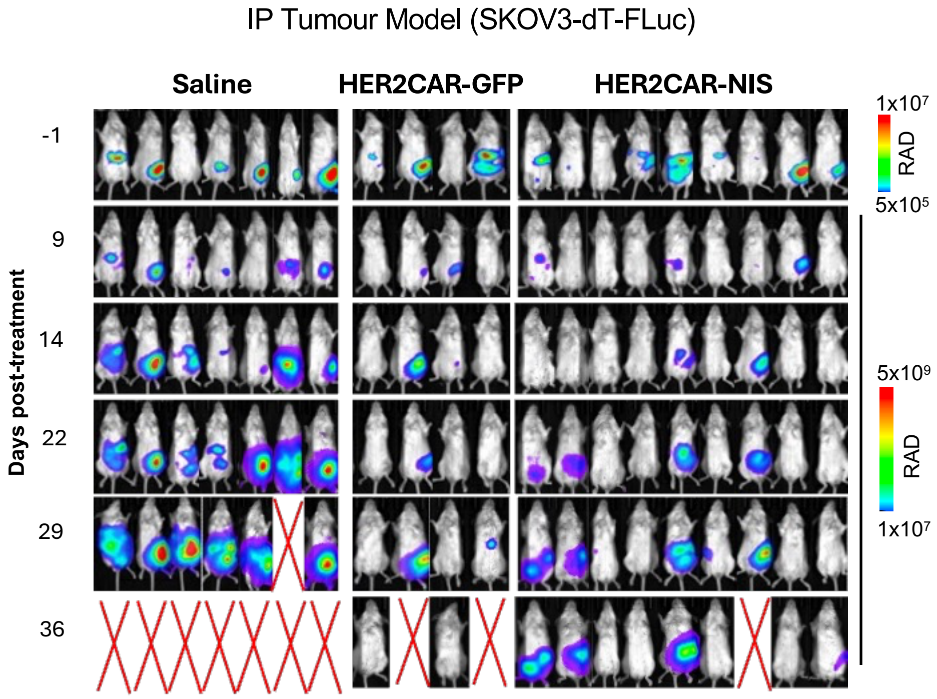


**Figure S4. BLI of intraperitoneal ovarian cancer tumours after HER2CAR-T cell therapy.**

Longitudinal BLI tracking of IP administered SKOV3-ip1 (dT+Fluc+) tumours before and after locoregional injections of saline, HER2CAR-GFP or HER2CAR-NIS T cells. Scale bars represent average radiance.
